# Increased alpha-helicity of a supercharged coiled-coil protein increases siRNA delivery efficiency of protein-lipid hybrid vehicle

**DOI:** 10.1101/2021.05.03.442303

**Authors:** Joseph Thomas, Julia Monkovic, Joseph A. Frezzo, Priya Katyal, Kamia Punia, Jin K. Montclare

## Abstract

Gene therapy has the potential to treat various diseases and has recently gained new interest due to the deployment nucleic acid based vaccines for COVID-19. Despite these developments, there still remains a need for further development of gene delivery vehicles to increase their safety and efficacy.. We have recently developed a lipoproteoplex (LPP) consisting of a super-charged coiled-coil protein (CSP) and a cationic liposomal carrier, that has the ability to condense nucleic acids and deliver them *in vivo*. The LPP is distinct from other liposomal gene delivery systems in that it utilizes a modular protein component to drive transfection activity as opposed to relying on the passive effects of the cationic lipids. A CSP library has been rationally designed to improve the efficacy of the LPP compared to the parent protein via improved alpha-helical structure and increased nucleic acid binding through the use of extended histidine tags and increased positive charge. The secondary structure and nucleic acid binding ability of each library member was assessed, then compared to functional transfection data in NIH-3T3 mouse fibroblasts. Structural and functional data suggests that increasing alpha-helicity of the protein component of the LPP compared to the parent sequence doubles nucleic acid binding affinity and increases transfection activity almost 3-fold with a favorable safety profile.

## 1. Introduction

Nucleic acids are a class of biomolecule with very high therapeutic potential, and have recently gained new attention due to their use in the COVID-19 vaccine [1–4]. Nucleic acids are vulnerable to degradation by nucleases in the body and suffer from low transfection efficiencies on their own, requiring the use of delivery vehicles to make them therapeutically viable [5–7]. The most commonly used gene delivery vectors currently are based on modified viruses and show high transfection ability [6, 8]. Since these vehicles are derived from viruses, the immune system can recognize and clear them from the body inhibiting their efficacy at high doses. Due to this, these vectors are primarily deployed in tissues such as the eye where the immune system has a very limited presence [9]. These delivery systems are also costly to produce, since they utilize complex manufacturing schemes [1, 10]. Non-viral vectors based on liposomes [11, 12], peptides [13–15], proteins [16–18], and polymers [19–21] have also been developed to address the safety concerns of viral vectors. These non-viral vectors are less immunologically active inside the body but have lower transfection efficiencies. This necessitates higher dosing to see a desired therapeutic effect, which often leads to an immune response in the body [10, 22]. There is a need to develop a modular platform technology that combines the high transfection efficiency of viral vectors with the favorable immune profile of non-viral vectors. This would allow for a wide range of nucleic acids such as plasmids, short interfering RNA (siRNA), and messenger RNA (mRNA) to be delivered safely anywhere in the body.

Synthetic biology and protein engineering techniques offer an attractive route to develop nextgeneration gene delivery vehicles since materials can be rapidly produced and screened [23, 24]. Protein-based materials can be rationally designed at the amino acid level to incorporate desired functionalities [25, 26] and have the benefit of being able to be produced recombinantly. Previous work has demonstrated that the secondary structure of proteins must be taken into account when designing nucleic acid binding materials [27]. In nature, the majority of nucleic acid binding proteins are alpha-helical and positively charged [28, 29]. This can allow for sequence specific interactions through the orientation of side chains able to form hydrogen bonds with specific nucleic acid base pairs [30], or sequence independent interactions through electrostatic interactions between positively charged amino acids and the negatively charged nucleic acid phosphate backbone [31]. The degree of alpha-helicity is also important, as more organized alpha-helices have been shown to form more energetically favorable interactions with nucleic acid binding partners [32]. This is facilitated by the insertion of a portion of the helix into the major or minor groove of the nucleic acid, which requires a high degree of secondary structure to occur [28, 32].

Our group has previously developed a protein-lipid hybrid vehicle known as a lipoproteoplex (LPP) that can safely deliver nucleic acids with a high efficiency [16, 17, 33]. The LPP utilizes a coiled-coil protein derived from the cartilage oligomeric matrix protein coiled coil (COMPcc) that self assembles into a homopentamer with a central hydrophobic pore [34]. Eight surface exposed residues of COMPcc have been mutated into positively charged arginine to create the coiled-coil supercharged protein, referred to as CSP8 [33]. CSP8 maintains the pentamer self-assembly and is able to bind to plasmid DNA and siRNA [17, 33]. When CSP8 is combined with a cationic liposome to create the LPP, this vehicle is able to transfect cells *in vitro* and has been used to improve the wound healing time of diabetic ulcers in a humanized mouse model through the delivery of an siRNA targeting Kelch-like ECH-associated protein 1 which acts a skey repressor of the Nrf2 pathway and plays an important role in wound antioxidant management [16, 35]. Our vehicle is unique from liposome-based delivery vehicles since it utilizes the functional protein component as the driver of transfection success. This allows for iterative improvement of our delivery platform via manipulation of the amino acid sequence of the protein component. Moreover, we can utilize an expansive design space that is not available to non-viral vehicles based on other material types.

To further improve the delivery efficacy of the LPP, a library of CSP8 mutants has been rationally designed to improve siRNA binding and delivery (**Figure 1**). Secondary structure of the mutants is assessed, and relative siRNA binding affinity is determined. Transfection activity and cytoxicity assays are carried out in mouse fibroblast cells, to compare functional data of the LPP to secondary structure and siRNA binding of each CSP8 mutant. CSP mutants with decreased alpha-helicity compared to the parent CSP8 show lower siRNA binding affinity and transfection activity. The variant, N8, has increased alpha-helicity compared to the parent CSP8 and has a higher siRNA binding affinity. The N8 LPP showed the highest transfection efficiency of the library as well, showing a clear relationship between protein secondary structure and siRNA binding/delivery.

**Figure 1:**
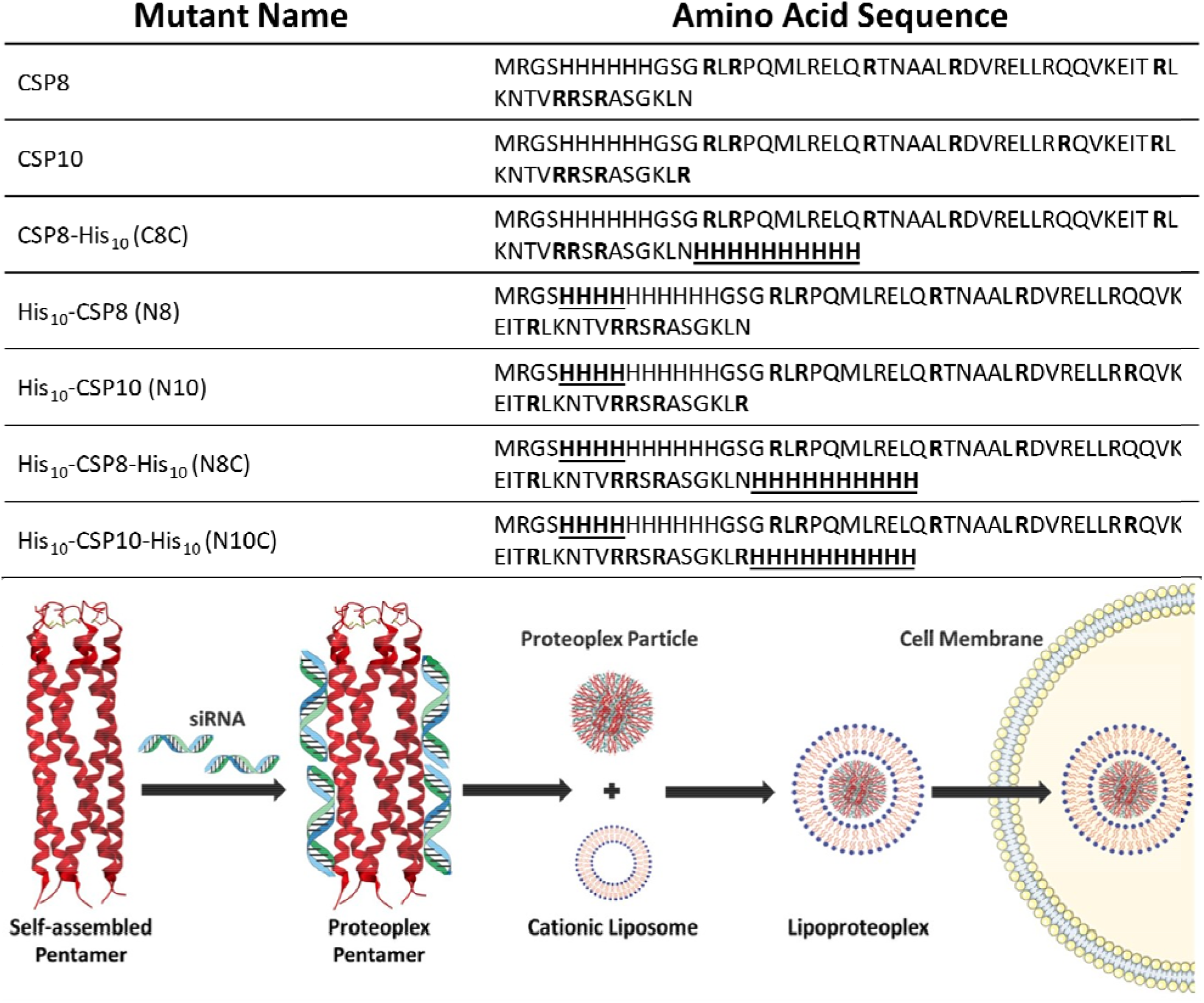
Amino-acid sequences of CSP8-based protein library (top). Mutated arginine residues are in bold and inserted histidine residues are in bold and underlined. Schematic of lipoproteoplex selfassembly (bottom). The supercharged protein variant binds to siRNA through charge-charge interactions. The protein-bound siRNA is then complexed with a cationic liposome to cross the cell membrane.

## 2. Materials and Methods

### 2.1 Materials

Primers were purchased from Eurofin MWG Operon (Huntsville, AL) and pfuUltraDNA polymerase was purchased from Stratagene (Santa Clara, CA) Q5 site-directed mutagenesis kit and DpnI restriction enzyme were purchased from New England Biolabs (Ipswich, MA). Phenylalanine auxotrophic Escherichia *coli* strain (AFIQ) were a gift from David Tirrell (California Institute of Technology). Tryptone, sodium chloride, yeast extract, tryptic soy agar, ampicillin, chloramphenicol, HisPur Ni-NTA agarose, sodium hydroxide (NaOH), isopropyl β-D-1-thiogalactopyranoside (IPTG), Tris hydrochloride (Tris HCl), imidazole, Pierce bicinchoninic acid (BCA) assay kit, Pierce snakeskin dialysis tubing 3.5 kD Molecular weight cut off (MWCO), urea, Coomassie Brilliant Blue R-250 dye, ethidium bromide, and sodium dodecyl sulfate (SDS) were acquired from Thermo Fisher Scientific. Hydrochloric acid (HCl) was purchased from VWR. Macrosep and microsep advance centrifugal 3K MWCO devices and 0.2 μm syringe filters were purchased from PALL. Acrylamide/bis solution (30%) 29:1 and natural polypeptide sodium dodecyl sulfate polyacrylamide gel electrophoresis (SDS-PAGE) standard were purchased from Bio-Rad. Agarose was purchased from Sigma-Aldrich. NIH-3T3 cells were obtained from Cell BioLabs. Dulbecco’s Modified Eagle Medium (DMEM), improved Minimal Essential Medium (OPTI-MEM), pen/strep, human insulin, L-glutamine, fetal bovine serum (FBS), non-essential amino acids (NEAA), PBS, Lipofectamine 2000, siRNA targeting green fluorescent protein (siGFP), T-75 flasks, and 24-well plates were purchased from Thermo Fisher.

### 2.2 Site Directed Mutagenesis and polymerase chain reaction (PCR) Assembly

The CSP8 gene in pQE30 vector was used as a template to perform multiple mutations. Two additional arginine residues were substituted into the parent CSP8 to produce CSP10 in a two-step sequential protocol and confirmed via sequencing. Gene sequences for the CSP library were generated using the Q5 site-directed mutagenesis kit. Site directed mutagenesis was performed using the manufacturer’s suggested protocol and the resulting sample was digested with DpnI enzyme for 3 hours at 37°C. The DpnI digested sample was transformed into XL-1 blue cells and insertions were confirmed via sequencing.

### 2.3 Protein Expression and Purification

Vectors from the CSP mutant library were transformed into AF-IQ *E. coli* cells and incubated on tryptic soy agar plates at 37°C to grow for 12-16 hours. The agar plates contained 0.2 μg/μL ampicillin and 0.034 μg/μL chloramphenicol for all protein variants. Single colonies were then selected following incubation and used to inoculate starter cultures. These starter cultures contained 5mL of lysogeny broth (LB) and 0.2 μg/μL ampicillin and 0.034 μg/μL chloramphenicol. The small culture tubes were incubated overnight at 37°C and 350rpm and were then used to inoculate the large-scale protein expression. Expression flasks containing 500 mL lysogeny broth with antibiotic treatment were incubated at 37°C and 350rpm for approximately 4 hours until an optical density (OD_600_) reading of ~0.8-1.0 was obtained. Following, the protein expression was induced using 0.2 μg/μL IPTG and incubated for another three hours at 37°C. Cells were then harvested via centrifugation at 5000 x g and 4°C for 45 minutes. Cell pellets were stored at −80°C until needed for purification.

Overexpression of each protein variant was confirmed via SDS-PAGE **(Figure S2)**. Cell pellets were thawed and resuspended into a buffer containing 50 mM Tris HCl, 100 mM NaCl, 20 mM imidazole, and 6M urea at pH 8.00. After resuspension, cells were lysed via sonication at 65% power, with 5 seconds on followed by 15 second off for a total on time of 1 minute and 30 seconds, for two full cycles. Following sonication, lysates were spun down to pellet cell debris and the lysate was incubated on Ni-NTA beads over night at 4°C on a rotisserie shaker (Labquake, Barnstead/Thermolyne). Purification was performed using a 10 mL gravity flow column (Thermo Fisher) at 4°C. Beads were washed with a buffer containing 50 mM Tris HCl, 100 mM NaCl, and 20 mM imidazole at pH 8.00. Mutants containing extended N-terminal histidine tags (N8, N10, N8C, N10C) also had 6 M urea in their wash buffer. Proteins were eluted from the beads using increasing amounts of imidazole (100 mM to 1M). Mutants containing extended N-terminal histidine tags utilized an imidazole range of 50 mM to 1M. Purity of proteins was confirmed via SDS-PAGE **(Figure S3)**. Proteins were dialyzed at room temperature to remove imidazole into a buffer containing 50 mM Tris HCl and 100 mM NaCl at pH 8.00. The C8C variant was found to aggregate in 100 mM NaCl, so was dialyzed at room temperature into a buffer containing 50 mM Tris HCl at pH 8.00 and NaCl was added immediately preceding experiments.

### 2.4 Circular Dichroism Spectroscopy

Protein was diluted to a concentration of 10 μM in buffer (50 mM Tris HCl, 100 mM NaCl, pH 8.00). Using the buffer as a blank, the protein was analyzed using the J-815 Circular Dichroism (CD) spectrophotometer with a 1 cm Quartz cuvette. Wavelength scans were taken from 200 nm to 250 nm with a 1 nm interval at 20°C to analyze secondary structure. Secondary structure was also analyzed as the protein was heated to 85°C and subsequently cooled back down to 20°C. Three wavelength scans were taken at each temperature, with an increase of 1°C/min in between temperatures. The observed ellipticity value (Θ) was converted into mean residue ellipticity (MRE) using the standard equation Θ_MRE_ = Θ/(10*cpl*) where *c* is the molar concentration of the protein, *p* is the path length in centimeters and *l* is the number of amino acids [33]. Melting behavior was analyzed by collecting the signal intensity at 222 nm over the temperature range of 20-85°C with a 1°C pitch. The fraction folded was derived using equation F = (Θ_A_ – Θ_U_)/(Θ_N_ – Θ_U_), where, Θ_A_ is the MRE observed at given temperature, Θ_U_ is MRE value for completely unfolded protein and Θ_N_ is the MRE value of completely folded protein that is considered at 20 °C [34]. The fraction folded curve was fit to a polynomial using GraphPad Prism software and used to calculate the melting temperature (T_m_). Percentage alpha-helicity was calculated using both a computational method and a method based on comparing the Θ_222_ to that of an ideal helix. The computational method utilized the Contin method using the CDPro package [36]. Spectra were input into the program and the alpha-helicity was determined using a reference set of 56 proteins. The fraction helicity compared to an ideal helix was calculated using the equation f = 100[(Θ)_222_/(Θ_max_)_222_] where (Θ_max_)_222_ = −40000[1-(2.5/*n*)] where *n* is the number of amino acid residues in the protein [34]. All data represented an average of three trials.

### 2.5 Electrophoretic Mobility Shift Assay

Non-coding control siRNA (Applied Biosystems Negative Control #1) was diluted to 10 μM in distilled water and the protein of interest was diluted in varying ratios with 50 mM Tris HCl, 100 mM NaCl, pH=8.00 buffer. 10 μL of siRNA was added to 10 μL of each protein dilution to form increasing weight/weight ratios ranging from 0.5:1 to 10:1. The reaction tubes were then left to incubate for 30 minutes at room temperature to allow the complexes to form. The resulting proteoplexes were loaded into a 2% agarose gel containing 200 μg/mL ethidium bromide, using a nucleotide ladder and naked siRNA as references for siRNA migration. The agarose gel was imaged under UV-light exposure immediately following electrophoresis. Full binding was determined to be the weight/weight ratio at which no free siRNA was observed in the gel. All binding data was completed in triplicate, and representative gel images are shown.

### 2.6 Transfection Assay

NIH-3T3 cells expressing GFP (ex/em 488/509) were maintained in a Symphony incubator (VWR, USA) at 37⍰C and 5% CO_2_ and grown in cell culture grade T-75 flasks. Cells were passaged when confluency reached 70-90%. Cells were grown in DMEM media supplemented with 10% FBS, 1% pen/strep, 0.1 mM NEAA, and 1% L-glutamine. For transfection studies, cells were seeded on 24-well plates at a density of 40k cells/well in full growth media and allowed to attach for 24 hours at 37⍰C and 5% CO_2_. At the end of this incubation, the full growth media was swapped for OPTI-MEM transfection media and transfection complexes delivering siGFP were added. Cells were exposed to the transfection complexes for 24 hours at 37⍰C and 5% CO_2_ before GFP fluorescence was read on a plate reader (SpectraMax, Molecular Devices, USA). Fluorescence signals were normalized to untreated cells.

Transfection complexes were prepared by mixing supercharged protein variant with siGFP (220 ng) at either a 10:1 or 3:1 protein/siGFP weight ratio in pure water for 30 minutes at room temperature. After this incubation, Lipofectamine 2000 (L2000) was diluted in OPTI-MEM according to the manufacturer’s specification and the protein-bound siGFP complexes were added dropwise. This L2000/protein-bound siGFP mixture was incubated at room temperature for 10 mins to form the LPP. The amount of L2000 used was such to result in a 4:1 weight ratio of L2000:siGFP. When the incubation was complete the LPP complexes were added dropwise to the NIH-3T3 cells for a final siGFP dose of 22 ng/well. As a positive control L2000 bound to siGFP was used and naked siGFP was used as a negative control. Data is presented as a mean of triplicates ± standard deviation.

### 2.7 Viability Assay

Cell viability 24 hours after treatment was measured using the CCK-8 kit (Dojindo Molecular Technologies, USA). Following fluorescence readings of the NIH-3T3 cells, fresh OPTI-MEM media was added to the plates alongside the CCK-8 reagent (10% of well volume) according to manufacturer’s specification. The cells were placed back in the incubator (37⍰C and 5% CO_2_) for one hour, and the absorbance at 450 nm was read. Cytotoxicity was determined by normalizing the absorbance signal to untreated cells. Data is presented as a mean of triplicates ± standard deviation.

### 2.8 Statistical Analysis

Statistics were computed for each appropriate experiment using the mean of a triplicate ± standard deviation. Data was compared using a Students t-test and a *p* value < 0.05 was used to determine statistical significance.

## 3. Results

### 3.1 Design Rationale

Mutants were designed to incorporate increased positive surface charge as well as extended histidine tags at both the N- and C-terminus. We hypothesize that increasing positive charge would lead to higher siRNA binding but may disrupt the secondary structure of the mutants. Since the LPP utilizes a coiled-coil supercharged protein, the alpha-helicity of the protein may be crucial for vehicle assembly and function. Extended histidine tags are incorporated to take advantage of the proton sponge effect during endocytosis of the LPP, which is hypothesized to lead to delivery of a higher amount of functional siRNA.

### 3.2 Protein Expression and Purification

PCR based mutation protocols were used to introduce extra positively charged arginine residues and extended histidine tags into the parent CSP8 sequence **(Figure S1).** All mutants were found to express in high amounts as determined by gel electrophoresis **(Figure S2).** CSP8 and CSP10 were both found to be readily soluble under native purification conditions, while mutants containing extended histidine tags were found to be more susceptible to aggregation and were therefore purified under denaturing conditions (6M urea) **(Figure S3).**

### 3.3 Protein Secondary Structure Analysis

Circular dichroism (CD) spectroscopy was used to determine the secondary structure of the supercharged library members. Previous work has shown that CSP8 is alpha-helical [33], but it was unknown how further mutations may affect protein structure and pentameric assembly. Wavelength scans revealed that all library members were alpha-helical **(Figure 2),** many of the library members were less alpha-helical than N8. Two of the mutants, N8 and C8C, were found to be more alpha-helical than CSP8. The Θ_222_/Θ_208_ was also examined as a measure of the coiled-coil assembly. All of the mutants had a ratio close to 1.00, indicating a coiled-coil pentamer assembly, except for N8C and N10C. These proteins had a ratio >1.0 indicative of higher order aggregation. This was confirmed during protein handling when the N8C and N10C mutants were prone to visible aggregation. Melting curves showed that all mutants could reversibly melt, spontaneously regaining alpha-helical structure upon cooling **(Figure S4).** Melting temperature calculations **(Figure S5)** revealed a T_m_ of 37.12 ± 9.97 ⍰C for CSP8. The addition of two positively charged arginine residues to yield CSP10 decreased the T_m_ to 32.12 ± 4.16 ⍰C. The addition of the extended N-terminal histidine tag to CSP8 to yield N8 increased the T_m_ to 51.48 ± 9.71 ⍰C. Adding two more arginine residues to N8 to make N10 showed no major change in T_m_ with N10 melting at 52.12 ± 11.49 ⍰C. Adding the C-terminal histidine tag to both N8 and N10 slightly increased the T_m_ with N8C and N10C melting at 61.59 ± 2.23 ⍰C and 53.86 ± 4.11 ⍰C respectively. The addition of the C-terminal histidine tag to CSP8 to create C8C also increased the T_m_ to 58.39 ± 1.84 ⍰C. These findings are in line with observations during protein purification where mutants containing extended histidine tags on either the N- or C-terminus were more prone to aggregation, and likely required more heat to melt due to the stability of histidine aggregates.

**Figure 2:**
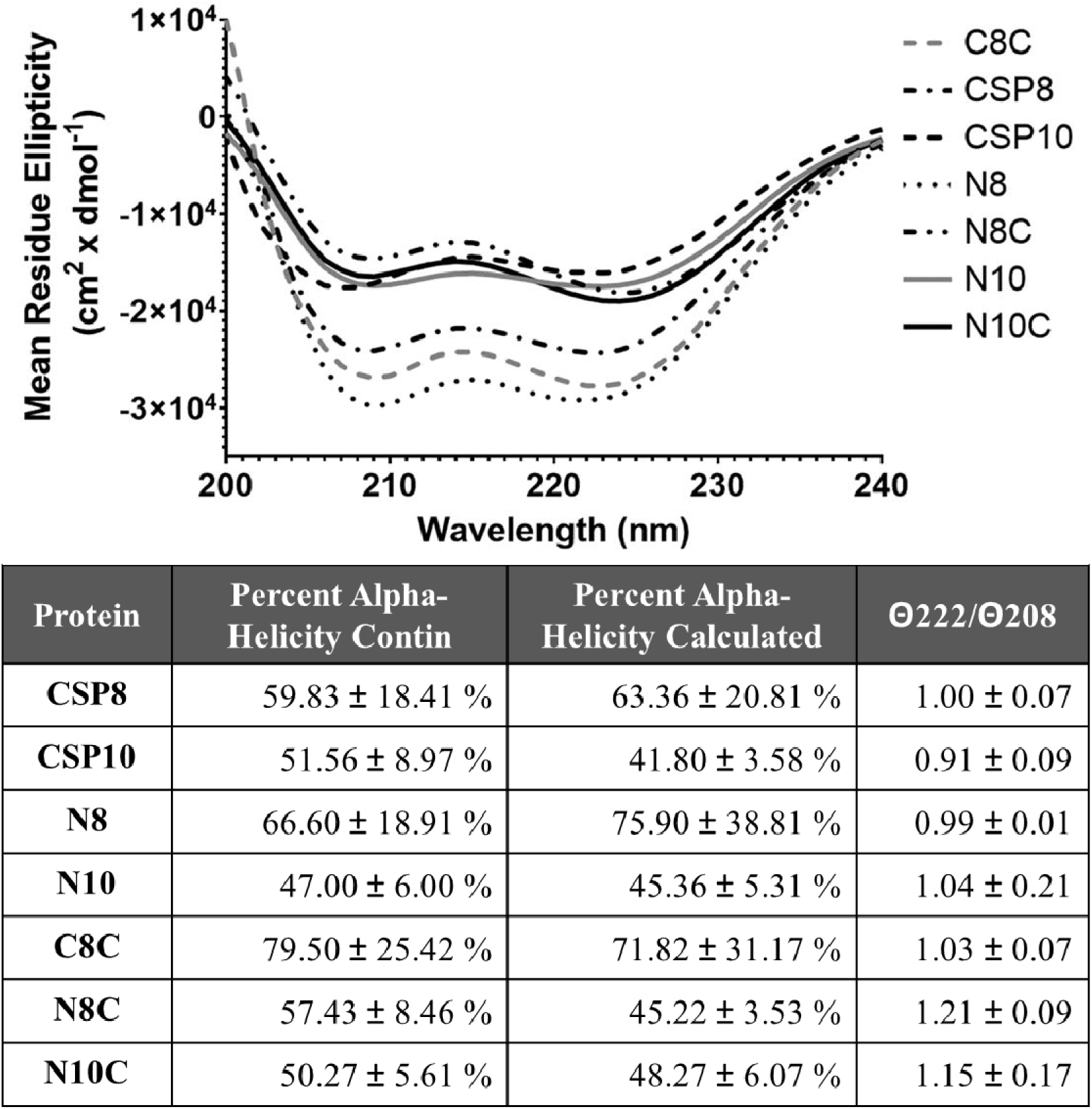
CD analysis of supercharged protein library. Wavelength scans (top) reveal that all proteins are alpha-helical. Structural calculations (bottom) reveal that proteins show varying degrees of alpha-helicity with N8 and C8C being the most alpha-helical.

### 3.4 Relative siRNA Binding Capacity

The ability of each protein mutant to complex with siRNA was assessed using an electrophoretic mobility shift assay (EMSA) **(Figure 3).** CSP8 was used as a baseline for binding activity since previous work showed that CSP8 can fully bind siRNA at a 2:1 weight ratio. N8 showed a higher binding activity and fully bound the siRNA at a 1:1 weight/weight ratio. CSP10 and N10 had similar binding activity to CSP8, fully binding at 2:1. N8C and N10C showed inhibited binding activity, only fully binding siRNA at a 5:1 and 3:1 ratio respectively. C8C had the lowest binding activity, and never reached binding saturation over the range tested.

**Figure 3:**
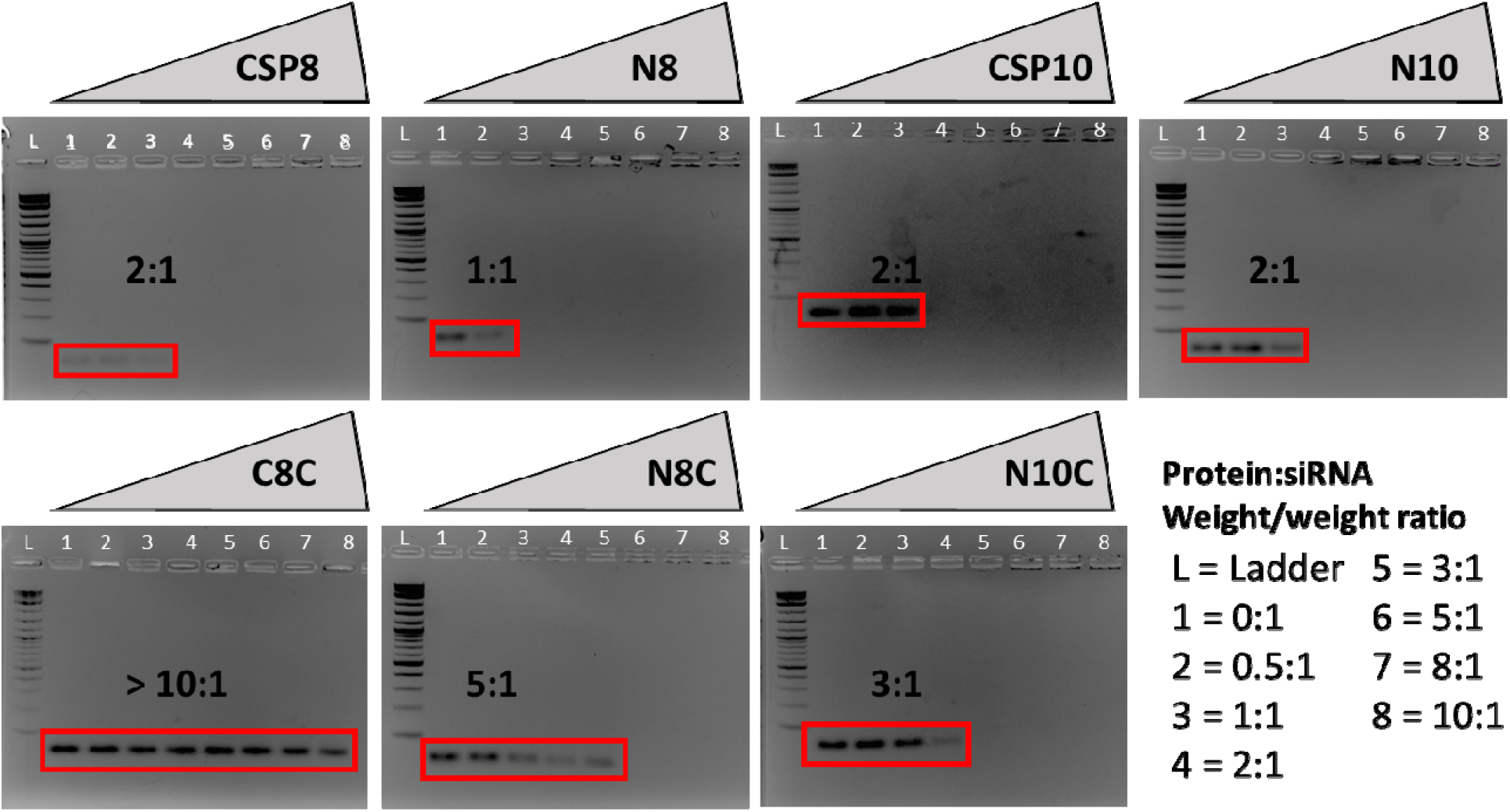
Relative binding capacities of protein mutants. A lower weight/weight binding ratio corresponds with higher binding activity. Protein bound siRNA remains in wells and free siRNA runs into gel (red boxes).

### 3.5 Transfection Activity of Lipoproteoplex Library

To determine the ability of each protein variant to influence the transfection efficiency of the lipoproteoplex (LPP), an siGFP knockdown assay was performed in NIH-3T3 GFP cells **(Figure 4).** This cell line exogenously expresses high levels of GFP regulated by a strong promoter. This makes it difficult to knockdown the GFP signal and makes the assay a suitable challenge for our LPP vehicle. The lipoproteoplex utilized commercially available Lipofectamine 2000 (L2000) as the cationic lipid component, as well as for a positive control without any added protein. L2000 loaded siGFP was able to knockdown the GFP signal 14.5 ± 2.4%. Naked siGFP was used as a negative control and demonstrated no significant knockdown. The LPPs were utilized to different protein:siGFP weight/weight ratios in order to see if the amount of protein played a role in knockdown efficiency. The L2000 amount was kept the same which resulted in final LPP formulations of either L2000:protein:siGFP 3:10:1 (LPP 10:1) or 3:3:1 (LPP 3:1). The CSP8 LPP 3:1 showed slightly higher knockdown than L2000:siGFP (18.5 ± 5.4%) but results did not reach statistical significance. None of the remaining LPP formulations outcompeted the L2000:siGFP group, except for the N8 LPP 3:1. The N8 LPP 3:1 showed a GFP knockdown of 46.8 ± 7.6%, significantly lower than the L2000 positive control (p=0.0017). even though most of the LPP formulations failed to outcompete L2000:siGFP, a trend is seen that the respective LPP 3:1 formulation shows a larger knockdown than the LPP 10:1 formulation.

**Figure 4:**
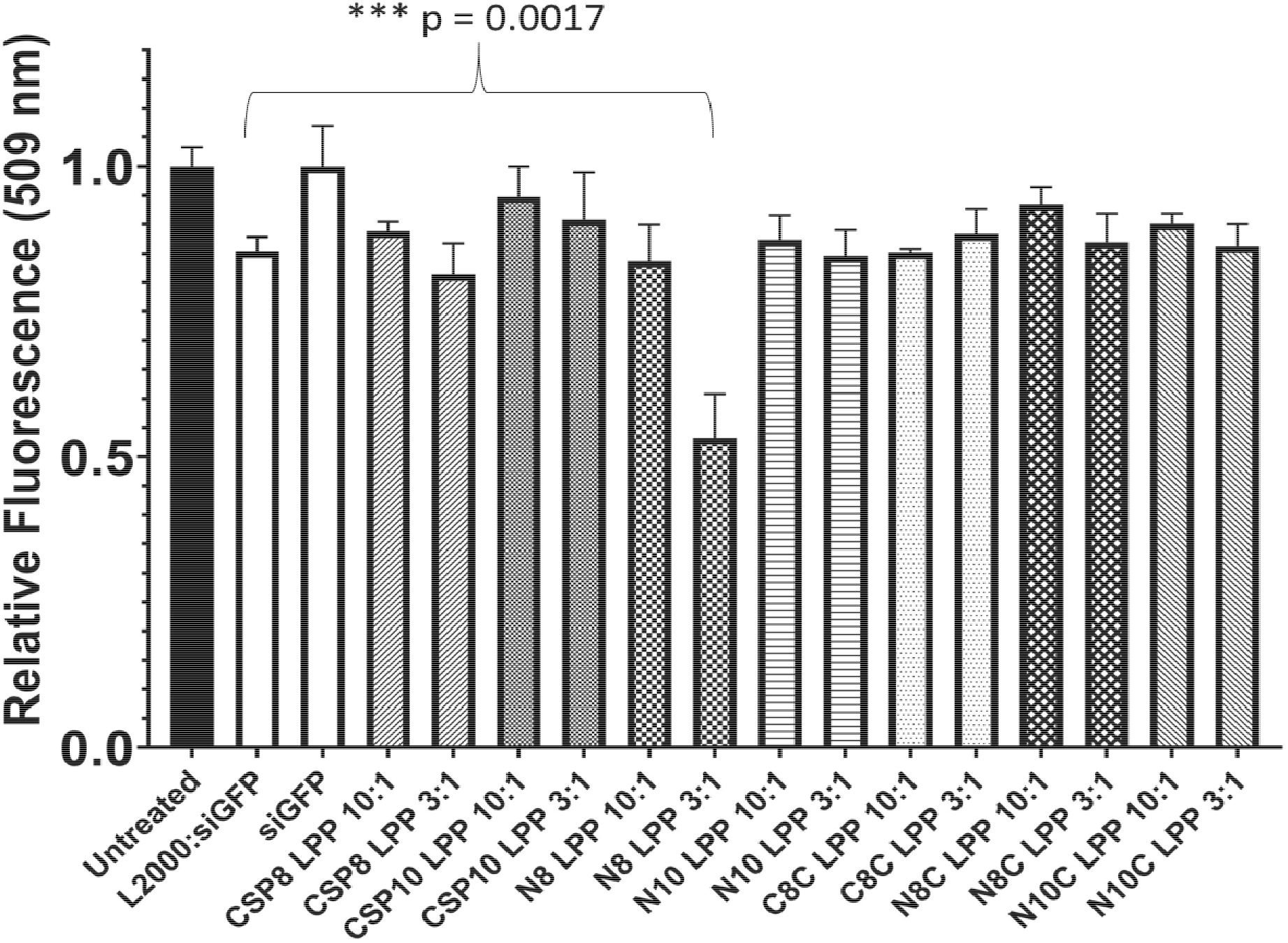
*In vitro* transfection efficiency of LPP formulations. Relative siGFP signal measured 24 hours post-transfection in NIH-3T3 GFP cells. The N8 LPP 3:1 shows the highest knockdown efficiency of all the treatment groups. All treatments are prepared using the same amount of siGFP and L2000 where appropriate. Fluorescence signals are normalized to untreated cells.

### 3.6 Cytotoxicity of Lipoproteoplex Library

Since the knockdown assay relied on a GFP readout from living cells, it was important to verify that knockdown effects were not coming from cell death due to vehicle toxicity. A CCK-8 assay was performed **(Figure 5)** immediately following GFP knockdown readings to correlate transfection activity with toxicity for each group. L2000:siGFP showed a relative cell viability of 84.8 ± 5.3%, which agreed with previously reported values and was not considered significantly toxic (viability < 80%) [16]. Naked siGFP showed a viability of 92.4 ±2.5%, which was not statistically different from the untreated cells. Viabilities for all LPP formulations tested did not significantly differ from the L2000:siGFP treatment, demonstrating that the addition of the supercharged protein component did not alter the safety profile of the LPP compared to the cationic L2000.

**Figure 5:**
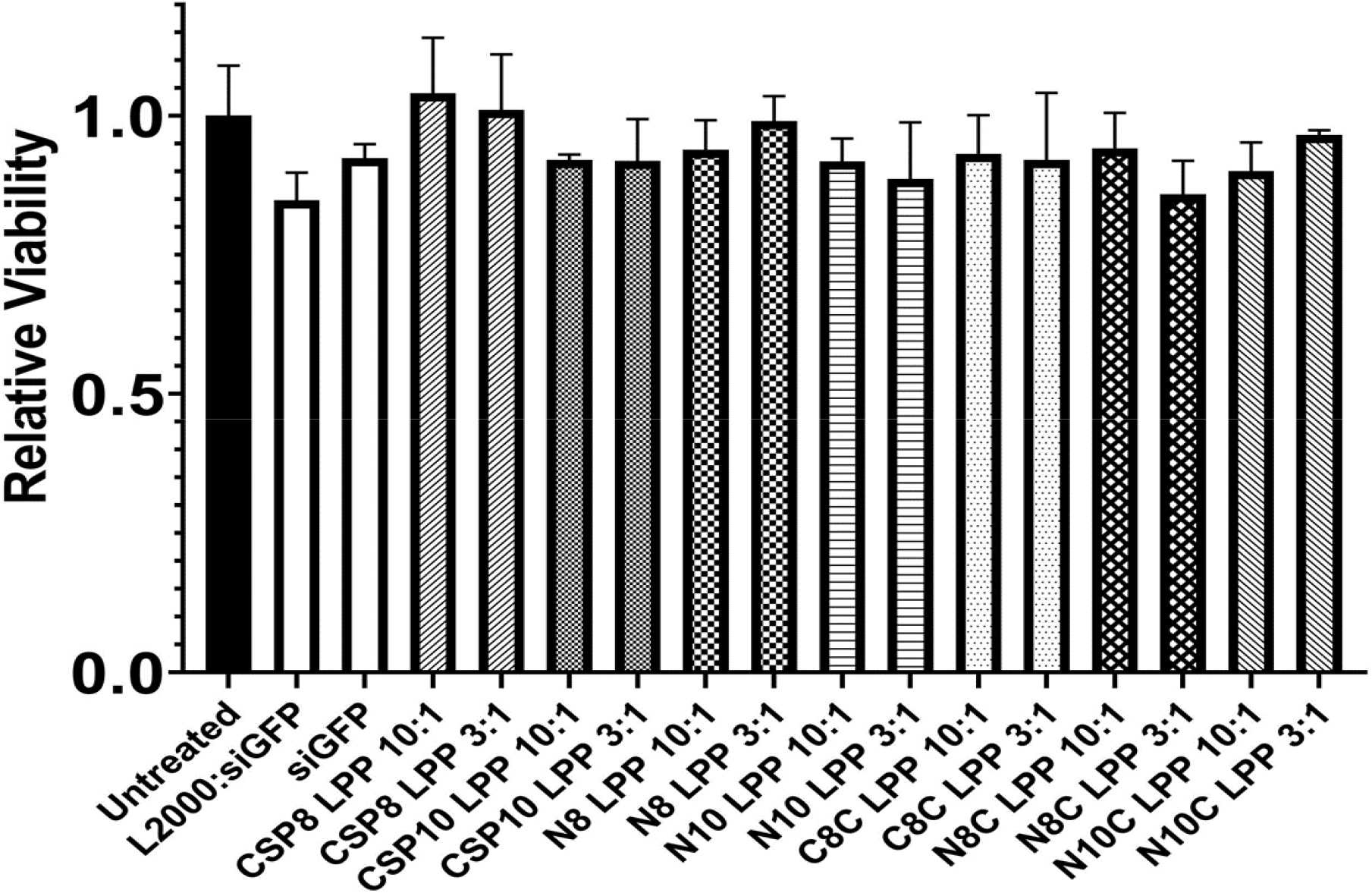
Cell viability measured 24 hours post-transfection via CCK-8 assay. Absorbance signals are normalized to untreated cells as a control (100% viability).

## 4. Discussion

The primary objective of this study was to create a more effective LPP via rational design of the supercharged coiled-coil protein component. Increased arginine residues and extended histidine tags were chosen in order to improve siRNA complexation and endosomal escape respectively. Interestingly, increasing positive charge (CSP8 vs. CSP10) did not improve siRNA complexation ability, but did have a negative effect on protein secondary structure. Comparing the transfection ability of CSP8 and CSP10 showed that the CSP10 LPP was not as efficient as the CSP8 LPP at GFP knockdown. The most striking difference in protein structure and function is observed when comparing CSP8 to N8. The addition of the extended N-terminal histidine tag (4 additional histidine residues) improved siRNA binding capacity, improved alpha-helicity, and when N8 is incorporated into the LPP it results in a vehicle with a very high knockdown efficiency. Combining the extended N-terminal histidine tag with increased arginine mutations in N10 unfortunately did not improve siRNA binding or transfection, and disrupted secondary structure compared to CSP8. It is clear from this data that the secondary structure of the protein component is a significant driver of siRNA binding and delivery efficiency. This is likely explained by previous work which showed that more ordered alpha-helices are better able to interact with the major and minor grooves of nucleic acids [28, 32]. The addition of the C-terminal histidine tag to CSP8 to generate C8C also greatly improved the alpha-helicity but abolished the siRNA binding ability. Due to this, the transfection activity of the C8C LPP was poor even though the secondary structure had been improved. The N8C and N10C variants also suffered from poor siRNA binding and transfection, likely due to the C-terminal histidine tags as well. These mutants were visibly more aggregated which could explain the poor siRNA binding. If the proteins form large aggregates, then they will have a decreased pool of surface accessible alpha-helices which can interact with the siRNA [32]. The presence of the extended N-terminal histidine tag appears to rescue binding ability somewhat, but the mutations have caused secondary structural disruptions compared to the parent CSP8.

General design principles for supercharged coiled-coils can be deduced from the behavior of the mutant library. Positive surface charge is crucial for electrostatic binding of the proteins to siRNA, but too many positively charged residues will begin to disrupt the structure as well as the nucleic acid binding which results in low transfection efficiency. Long N-terminal histidine tags (10 residues) exhibit a stabilizing effect on supercharged protein secondary structure and lead to an increased nucleic acid binding ability and a much higher transfection ability. C-terminal histidine tags appear to inhibit siRNA binding and result in non-functional protein variants. The final transfection ability of the mutants appeared to be a sum of the mutations to the protein, demonstrating a “less is more” principle in the design process since seemingly synergistic mutations could actually compete against each other and not yield the desired result. Future designs should aim to further improve protein secondary structure to create even more favorable protein-siRNA interactions [30], while maintaining a favorable positive charge density [31].

These findings are important as design principles need to maximize functional siRNA delivery while minimizing toxicity. The siGFP knockdown experiments demonstrated that cationic liposomal vehicles (L2000) are not efficient as they could be at siRNA delivery, and the addition of a functional protein component opens an entire design paradigm. The N8 LPP had almost a three-fold increase in transfection efficiency compared to L2000 alone as well as the CSP8 LPP with no significant increase in toxicity. The alpha-helicity of our supercharged protein can likely be improved even further, which would be hypothesized to create a next-generation lipoproteoplex even more effective than the current ones.

## 5. Conclusion

We have rationally designed a library of supercharged coiled-coil proteins to use in conjunction with cationic liposomes for efficient non-viral siRNA delivery. Our work has resulted in the protein N8 which outcompetes commercially available cationic liposomes as well as our previous protein CSP8. We have discovered that the secondary structure of N8 is key in dictating siRNA binding and ultimately transfection success. This echoes finding of other groups which indicate that alpha-helical structure is a prerequisite for protein-nucleic acid interactions. The next generation N8 LPP is able to improve transfection efficiency three-fold compared to the previous LPP without increasing cytotoxicity. Further studies are already underway to further optimize alpha-helicity and to uncover how protein secondary structure influences LPP assembly and internalization. Studies are also underway to optimize the lipid component of the LPP to further improve safety and efficacy.

## Acknowledgements

This work was supported by the National Science Foundation award number DMREF-1728858.

## References

1. Goswami, R., et al., Gene Therapy Leaves a Vicious Cycle. Front Oncol, 2019. 9: p. 297.

2. Wang, F., et al., Clinical translation of gene medicine. J Gene Med, 2019: p. e3108.

3. de Queiroz, N.M.G.P., et al., Vaccines for COVID-19: perspectives from nucleic acid vaccines to BCG as delivery vector system. Microbes Infect, 2020. 22(10): p. 515–524.

4. Pushparajah, D., et al., Advances in gene-based vaccine platforms to address the COVID-19 pandemic. Adv Drug Deliv Rev, 2021. 170: p. 113–141.

5. Hayat, S.M.G., et al., Gene Delivery Using Lipoplexes and Polyplexes: Principles, Limitations and Solutions. Crit Rev Eukaryot Gene Expr, 2019. 29(1): p. 29–36.

6. Chen, Y.H., M.S. Keiser, and B.L. Davidson, Viral Vectors for Gene Transfer. Curr Protoc Mouse Biol, 2018. 8(4): p. e58.

7. Viola, J.R., et al., Non-viral nanovectors for gene delivery: factors that govern successful therapeutics. Expert Opin Drug Deliv, 2010. 7(6): p. 721–35.

8. Schaffer, D.V., J.T. Koerber, and K.I. Lim, Molecular engineering of viral gene delivery vehicles. Annu Rev Biomed Eng, 2008. 10: p. 169–94.

9. Campbell, J.P., T.J. McFarland, and J.T. Stout, Ocular Gene Therapy. Dev Ophthalmol, 2016. 55: p. 317–21.

10. Riley, M.K. and W. Vermerris, Recent Advances in Nanomaterials for Gene Delivery-A Review. Nanomaterials (Basel), 2017. 7(5).

11. Akbarzadeh, A., et al., Liposome: classification, preparation, and applications. Nanoscale Res Lett, 2013. 8(1): p. 102.

12. Alshehri, A., A. Grabowska, and S. Stolnik, Pathways of cellular internalisation of liposomes delivered siRNA and effects on siRNA engagement with target mRNA and silencing in cancer cells. Sci Rep, 2018. 8(1): p. 3748.

13. Taylor, R.E. and M. Zahid, Cell Penetrating Peptides, Novel Vectors for Gene Therapy. Pharmaceutics, 2020. 12(3).

14. Baumhover, N.J., et al., Structure-Activity Relationship of PEGylated Polylysine Peptides as Scavenger Receptor Inhibitors for Non-Viral Gene Delivery. Mol Pharm, 2015. 12(12): p. 4321–8.

15. Bang, E.K., et al., Amphiphilic small peptides for delivery of plasmid DNAs and siRNAs. Chem Biol Drug Des, 2018. 91(2): p. 575–587.

16. Rabbani, P.S., et al., Novel lipoproteoplex delivers Keap1 siRNA based gene therapy to accelerate diabetic wound healing. Biomaterials, 2017. 132: p. 1–15.

17. Liu, C.F., et al., Efficient Dual siRNA and Drug Delivery Using Engineered Lipoproteoplexes. Biomacromolecules, 2017. 18(9): p. 2688–2698.

18. Numata, K., et al., Bioengineered silk protein-based gene delivery systems. Biomaterials, 2009. 30(29): p. 5775–84.

19. Alinejad-Mofrad, E., et al., Evaluation and comparison of cytotoxicity, genotoxicity, and apoptotic effects of poly-l-lysine/plasmid DNA micro- and nanoparticles. Hum Exp Toxicol, 2019. 38(8): p. 983–991.

20. Yang, S. and S. May, Release of cationic polymer-DNA complexes from the endosome: A theoretical investigation of the proton sponge hypothesis. J Chem Phys, 2008. 129(18): p. 185105.

21. Golan, M., V. Feinshtein, and A. David, Conjugates of HA2 with octaarginine-grafted HPMA copolymer offer effective siRNA delivery and gene silencing in cancer cells. Eur J Pharm Biopharm, 2016. 109: p. 103–112.

22. Tros de Ilarduya, C., Y. Sun, and N. Düzgüneş, Gene delivery by lipoplexes and polyplexes. Eur J Pharm Sci, 2010. 40(3): p. 159–70.

23. Wilson, C.J., Rational protein design: developing next-generation biological therapeutics and nanobiotechnological tools. Wiley Interdiscip Rev Nanomed Nanobiotechnol, 2015. 7(3): p. 330–41.

24. Bloom, J.D., et al., Evolving strategies for enzyme engineering. Curr Opin Struct Biol, 2005. 15(4): p. 447–52.

25. Hill, L.K., et al., Protein-Engineered Nanoscale Micelles for Dynamic. ACS Nano, 2019. 13(3): p. 2969–2985.

26. Yin, L., et al., Engineered Coiled-Coil Protein for Delivery of Inverse Agonist for Osteoarthritis. Biomacromolecules, 2018. 19(5): p. 1614–1624.

27. Lin, M. and J.T. Guo, New insights into protein-DNA binding specificity from hydrogen bond based comparative study. Nucleic Acids Res, 2019. 47(21): p. 11103–11113.

28. Rohs, R., et al., Origins of specificity in protein-DNA recognition. Annu Rev Biochem, 2010. 79: p. 233–69.

29. Garvie, C.W. and C. Wolberger, Recognition of specific DNA sequences. Mol Cell, 2001. 8(5): p. 937–46.

30. Steitz, T.A., Structural studies of protein-nucleic acid interaction: the sources of sequence-specific binding. Q Rev Biophys, 1990. 23(3): p. 205–80.

31. Yu, B., B.M. Pettitt, and J. Iwahara, Dynamics of Ionic Interactions at Protein-Nucleic Acid Interfaces. Acc Chem Res, 2020. 53(9): p. 1802–1810.

32. Rathnayake, P.V., et al., Trends in the Binding of Cell Penetrating Peptides to siRNA: A Molecular Docking Study. J Biophys, 2017. 2017: p. 1059216.

33. More, H.T., et al., Gene delivery from supercharged coiled-coil protein and cationic lipid hybrid complex. Biomaterials, 2014. 35(25): p. 7188–93.

34. Gunasekar, S.K., et al., N-terminal aliphatic residues dictate the structure, stability, assembly, and small molecule binding of the coiled-coil region of cartilage oligomeric matrix protein. Biochemistry, 2009. 48(36): p. 8559–67.

35. Soares, M.A., et al., Restoration of Nrf2 Signaling Normalizes the Regenerative Niche. Diabetes, 2016. 65(3): p. 633–46.

36. Sreerama, N., S.Y. Venyaminov, and R.W. Woody, Estimation of protein secondary structure from circular dichroism spectra: inclusion of denatured proteins with native proteins in the analysis. Anal Biochem, 2000. 287(2): p. 243–51.

