## Supplemental figures for "Increased alpha-helicity of a supercharged coiled-coil protein increases siRNA delivery efficiency of protein-lipid hybrid vehicle"

**Supplementary Materials**

CSP10 was created using CSP8 as a template by a polymerase chain reaction (PCR) in a 50 μL volume using high fidelity PfuUltra HotStart Taq polymerase (2.5 units/μL, 1 μL), 10x PfuUltra reaction buffer (5 μL), deoxynucleotide triphosphates mix (10 mM, 1 μL), plasmid DNA (200-300 ng/μL, 1 μL), diH_2_O (40 μL) and the following primers (125 ng/μL, 1 μL each) from MWG:

Q39R Fwd 5’-CGTGAACTGCTGCGTCGGCAGGTTAAAGAAATCAC-3’

Q39R Rev 5’-GTGATTTCTTTAACCTGCCGACGCAGCAGTTCACG -3’

N61R Fwd 5’-CGTCTGGTAAGCTTAGATAGCTGAGCTTGG-3’

N61R Rev 5’-CCAAGCTCAGCTATCTAAGCTTACCAGACG-3’

All other mutants were prepared with a Q5 site-directed mutagenesis kit using either CSP8 or CSP10 as a template. Single 12.5 μL reactions were prepared with Q5 Hot Start High-Fidelity 2X Master Mix (6.25 μL), 10 μM forward primer (0.625 μL), 10 μM reverse primer (0.625 μL), 25 ng/μL template DNA (1 μL) and nuclease-free water (4 μL).

His_10_-CSP8 (N8) and His_10_-CSP10 (N10) were prepared from CSP8 or CSP10 respectively using the following primers:

Fwd 5’-CACCATCACCATCACGGATCCGGT-3’

Rev 5’-ATGGTGATGGTGATGCGATCCTCTC-3’

To prepare CSP8-His_10_ (C8C), the following primers were used with CSP8 as a template:

Fwd 5’-ATGGTGATGGTGATGATTAAGCTTACCAGACGC-3’

Rev 5’-CACCATCACCATCACTAGCTGAGCTTGGACTCC-3’

His_10_-CSP8-His_10_ (N8C) and His_10_-CSP10-His_10_ (N10C) were prepared from CSP8 or CSP10 respectively using the following primers:

Fwd 5’-CACCATCACCATCACTAGCTGAGCTTGGACTCC-3’

Rev 5’-ATGGTGATGGTGATGTCTAAGCTTACCAGACGC-3’

**Figure S1:** Mutagenesis protocol and primer list for performed mutations.

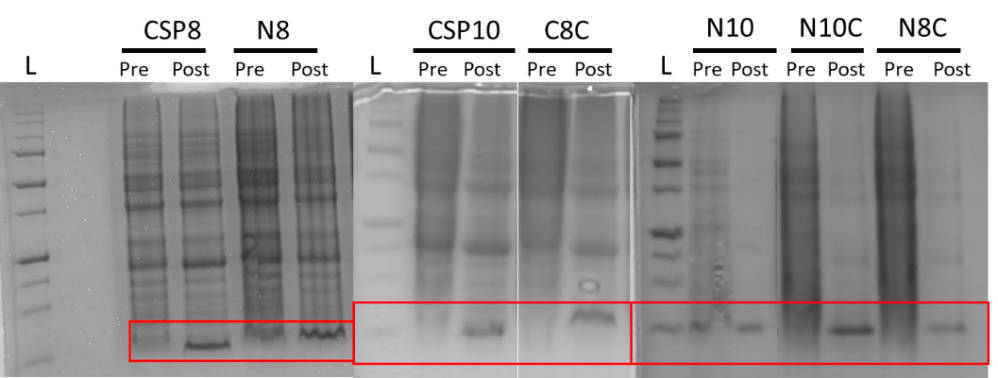

**Figure S2:** Expression gels for mutant library. Overexpression is indicated by a thick protein band around 7.5 kDa in the lanes labelled “post”.

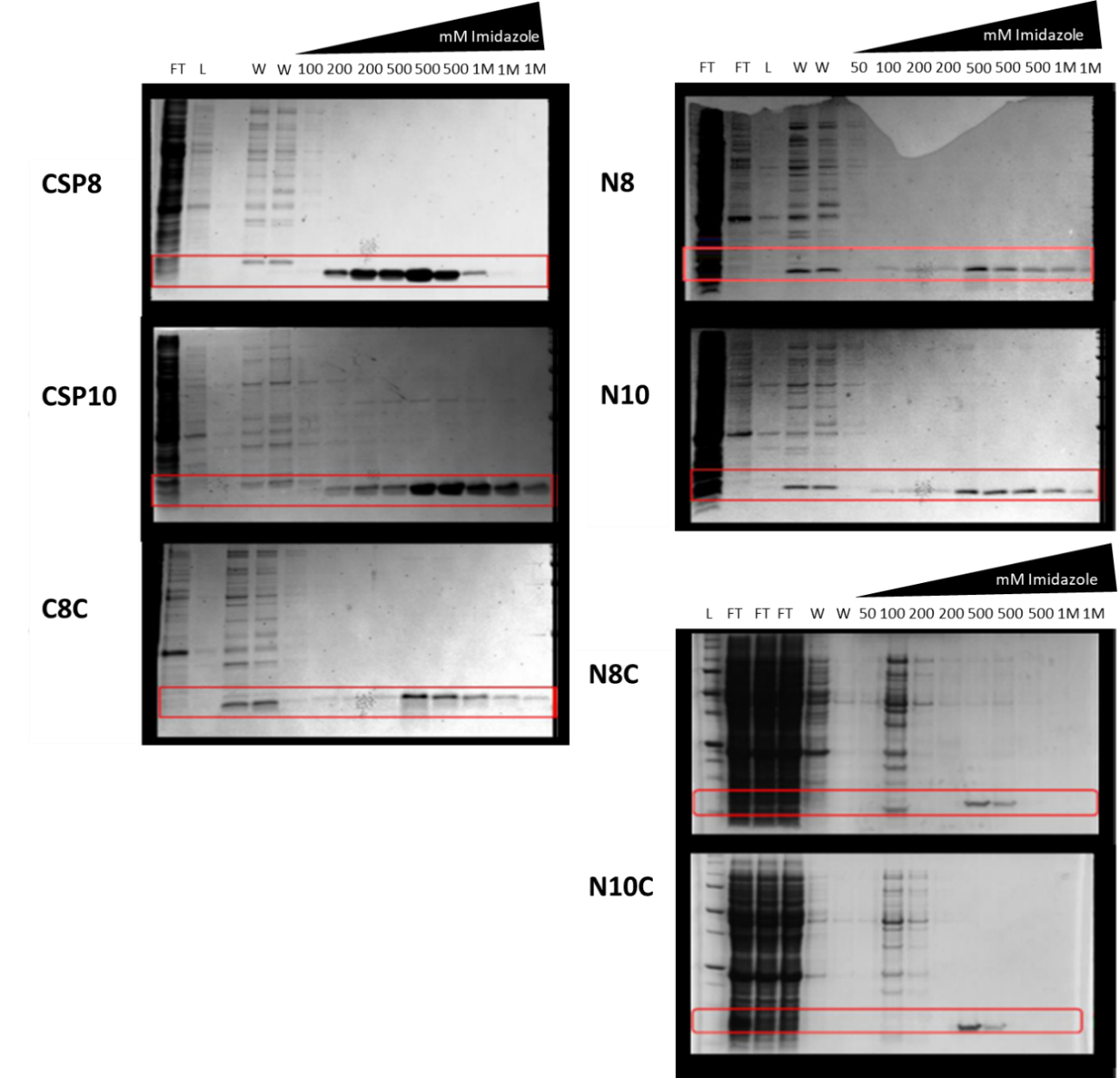

**Figure S3:** Purification gels for mutant library. Red boxes indicate mutant protein position in gel.

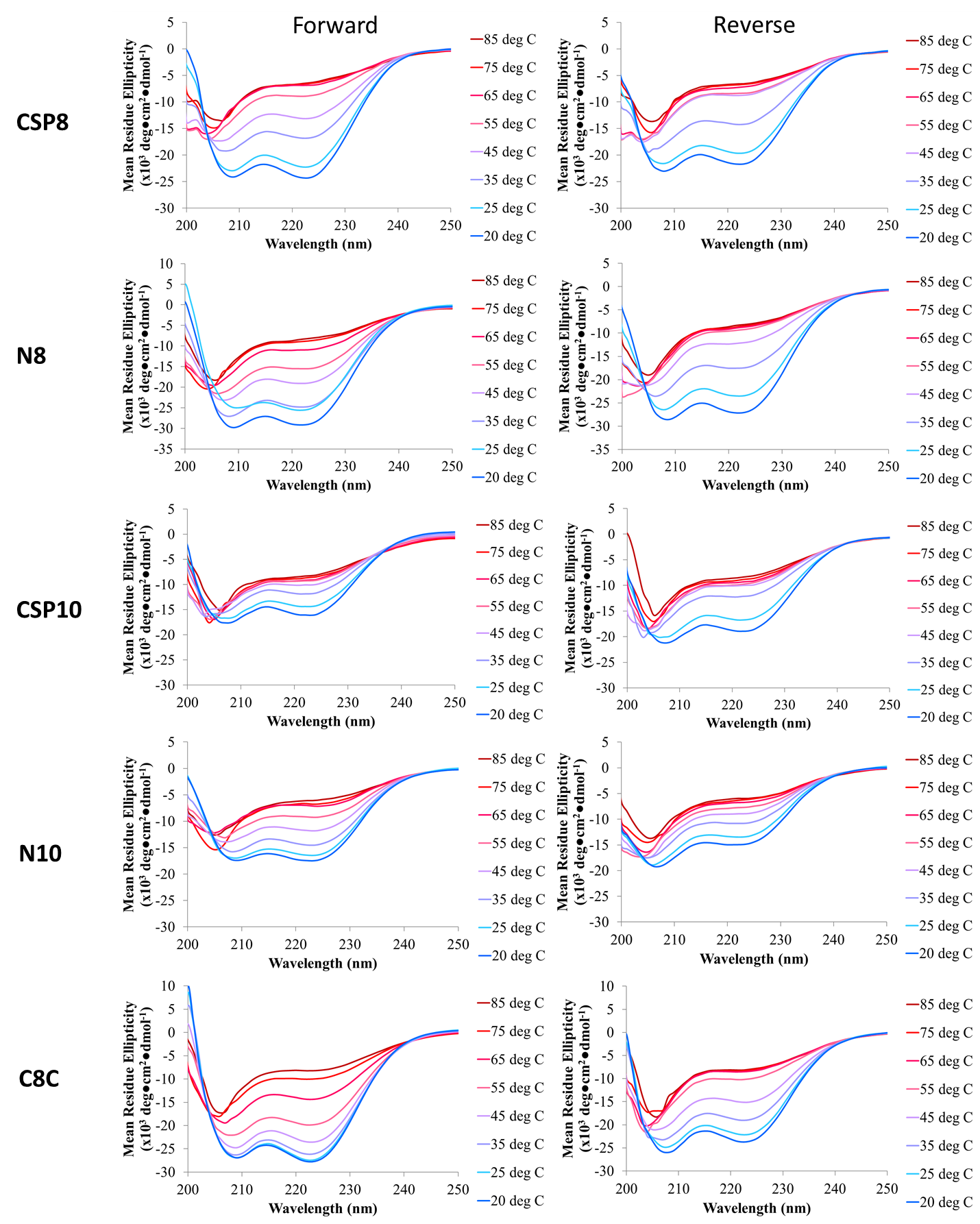

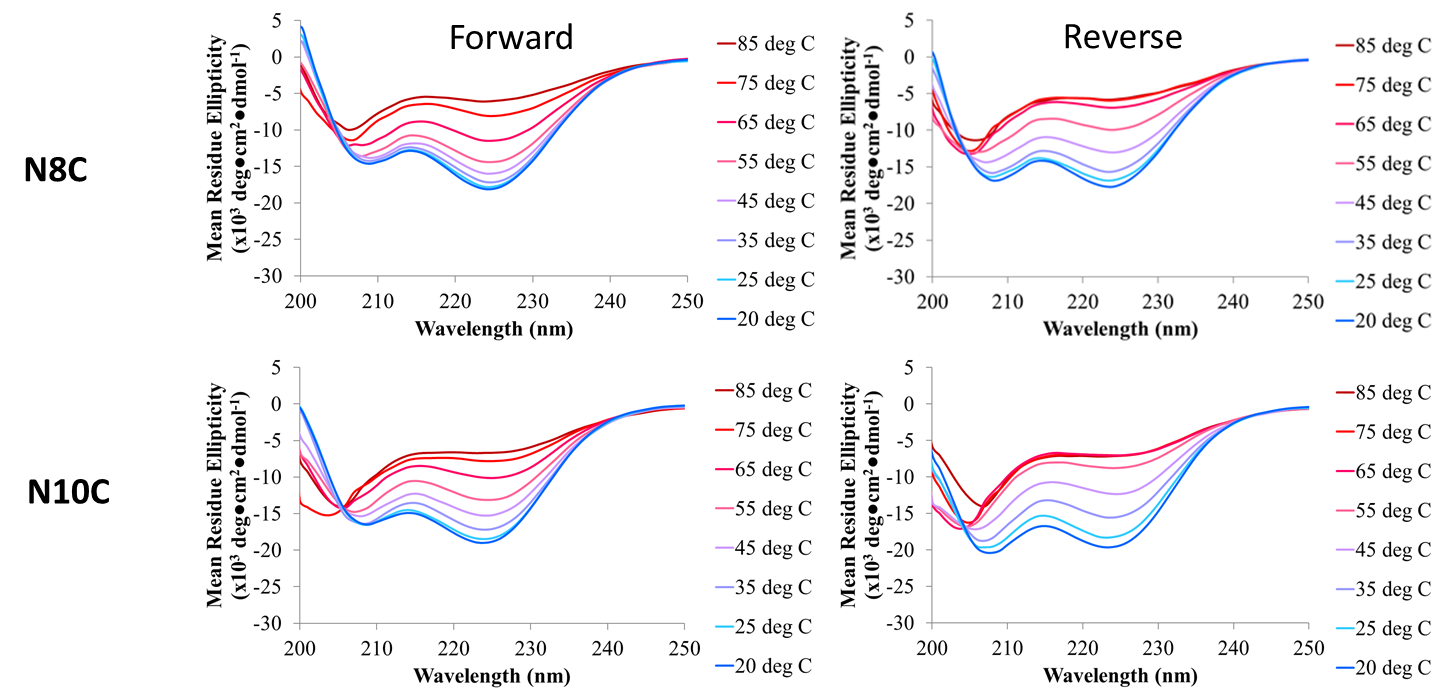

**Figure S4:** CD melt curves for forward (left) and reverse (right) experiments. Forward scans involve heating sample from 20ᵒC to 85ᵒC, and reverse scans involve cooling sample from 85ᵒC to 20ᵒC.

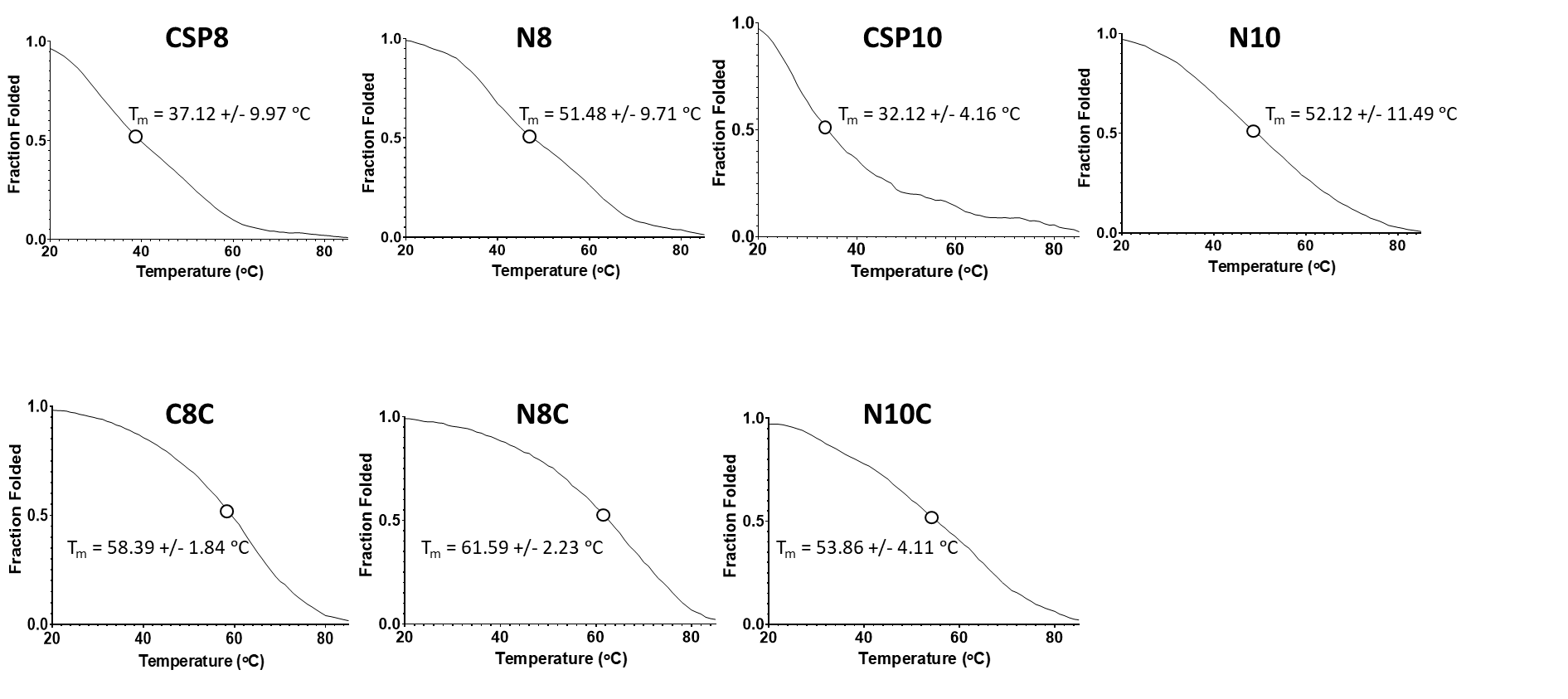

**Figure S5:** Fraction folded curves and melting temperatures (T_m_).
